# Tandem repeats contribute to coding sequence variation in bumblebees (Hymenoptera: Apidae)

**DOI:** 10.1101/201459

**Authors:** Xiaomeng Zhao, Long Su, Sarah Schaack, Ben M. Sadd, Cheng Sun

**Author notes:** Contributed equally (Xiaomeng Zhao, Long Su). Author for Correspondence: Cheng Sun, Key Laboratory of Pollinating Insect Biology of the Ministry of Agriculture, Institute of Apicultural Research, Chinese Academy of Agricultural Sciences, Beijing 100093, China.

## Abstract

Tandem repeats (TRs) are highly dynamic regions of the genome. Mutations at these loci represent a significant source of genetic variation and can facilitate rapid adaptation. Bumblebees are important pollinating insects occupying a wide range of habitats. However, to date, molecular mechanisms underlying the potential adaptation of bumblebees to diverse habitats are largely unknown. In the present study, we investigate how TRs contribute to genetic variation in bumblebees, thus potentially facilitating adaptation. We identified 26,595 TRs in the buff-tailed bumblebee (*Bombus terrestris*) genome, 66.7% of which reside in genic regions. We also compared TRs found in *B. terrestris* with those present in the whole genome sequence of a congener, *B. impatiens*. We found that a total of 1,137 TRs were variable in length between the two sequenced bumblebee species, and further analysis reveals that 101 of them are located within coding regions. The 101 TRs were responsible for coding sequence variation and corresponded to protein sequence length variation between the two bumblebee species. The variability of identified TRs in coding regions between bumblebees was confirmed by PCR amplification of a subset of loci. Functional classification of bumblebee genes where coding sequences include variable-length TRs suggests that a majority of these genes are related to transcriptional regulation. Our results show that TRs contribute to coding sequence variation in bumblebees and TRs may facilitate the adaptation of bumblebees through diversifying proteins involved in controlling gene expression.

## INTRODUCTION

Tandem repeats (TRs) are DNA tracts in which a short DNA sequence, dubbed a repeat unit, is repeated several times in tandem, and they are ubiquitous in the genomes of diverse species (Gemayel et al. 2010; Melters et al. 2013; Vinces et al. 2009). Most mutations in TRs are due to the variation in repeat unit number that occurs when one or more repeat units are added or deleted via a variety of different mutational mechanisms (e.g., DNA polymerase slippage (Tachida and Iizuka 1992)). Because they are known to be highly variable, TRs are also known as VNTRs (variable number of tandem repeats; Gemayel et al. 2010). A number of cellular processes (for example, DNA replication, recombination, faulty DNA damage repair, and other aspects of DNA metabolism) and external factors are known to cause mutations in TRs, thus the frequency of mutations at these loci is thought to be 100 to 10,000 times higher than point mutations (López et al. 2010; Paques et al. 1998; Rando and Verstrepen 2007; Schmidt and Mitter 2004; Tachida and Iizuka 1992).

Mutations in TRs can have phenotypic consequences. Firstly, mutations in TRs residing in coding regions can impact the structure, function, or processing of messenger RNAs or proteins. Several neurodegenerative diseases have been linked to the repeat unit number variation of TRs located in coding regions, the most famous case being the abnormal expansion of a CAG repeat in exon 1 of the *IT15* gene leading to Huntington’s disease (HD). Repeat numbers ranging from 6 to 35 are found in healthy individuals, whereas alleles with 40 repeats or more cause HD (Duyao et al. 1993; Gatchel and Zoghbi 2005).

In addition to their role in disease, TRs in coding regions also confer phenotypic variability without major fitness costs (reviewed in Kashi and King 2006). The repeat unit number variation in TRs located in *FLO1* gene in *Saccharomyces cerevisiae*generates plasticity in cell adhesion to substrates (Verstrepen et al. 2005). In canines, variable TRs located in developmental genes confer variability to skeletal morphology (Fondon and Garner 2004). Further, mutations in TRs located in non-coding regions can also have significant effects. Variable length TRs have been shown to influence transcription factor binding, as well as potentially changing DNA structure, packaging, and spatial dynamics, in addition to changing the secondary structure of RNA molecules once transcribed. Tandem repeats in promoters change gene expression in yeast (Vinces et al. 2009), and contribute to gene expression variation in humans (Gymrek et al. 2016). Therefore, given that TRs are highly mutable regions in the genome, and thus represent a significant source of variation, in cases where this variation is at loci influencing morphological, physiological and behavioral traits, it could facilitate adaptation to different environments (Feliciello et al. 2015; Fonville et al. 2011; Gemayel et al. 2010; Fidalgo et al. 2006; Tautz et al. 1986; Verstrepen et al. 2005; Vinces et al. 2009; Xu et al. 2017; Zhou et al. 2014).

Bumblebees (Hymenoptera: Apidae) are a genus of pollinating insects that play an important role in crop production and natural ecosystem services (Fontaine et al. 2006; Garibaldi et al. 2013; Velthuis and van Doorn 2006). They are distributed widely across the globe, from Greenland to the Amazon Basin and from sea level to altitudes of 5800 m in the Himalayas (Williams 1985). Bumblebees occupy a remarkably wide diversity of habitats, from alpine meadows to lowland tropical forest (Sakagami 1976). However, to date, molecular mechanisms underlying the adaptation of bumblebees to such a diverse array of habitats are largely unknown. Genetic variation is important for adaptation to new environments (Barrett and Schluter 2008; Lande and Shannon 1996; Paaby and Rockman 2014), however, little is known about sources or levels of genetic variation in bumblebees (but see (Lozier et al. 2011; Maebe et al. 2016)).

In the present study, we performed a systematic examination of TRs in the bumblebee genome and investigate their contribution to genetic variation in bumblebees. We further examine the functional significance of the genetic variation introduced by TRs to bumblebee genes. Lastly, we discuss the potential significance of the added genetic variation, especially as it may influence the regulation of gene expression.

## Materials and Methods

### Genomic sequences, annotation and predicted proteins

The genomic sequences, genome annotation, and predicted protein sequences of *Bombus terrestris* were downloaded from GenBank (https://www.ncbi.nlm.nih.gov/genome/2739, last accessed on April 5, 2016; GenBank assembly accession of GCF_000214255.1 [Bter_1.0]). The genomic sequences and predicted protein sequences of *Bombus impatiens* were downloaded from GenBank (http://www.ncbi.nlm.nih.gov/genome/3415, last accessed on April 5, 2016; GenBank assembly accession of GCA_000188095.2 [BIMP_2.0]).

### Bumblebee genomic DNA

The three worker specimens of *Bombus terrestris* were collected in the summer of 2017 in Burqin County, Xinjiang Uygur Autonomous Region, China from three different sites all within a 6-kilometer range of a previously collected conspecific (GPS coordinates: latitude 48.19179; longitude 87.02355). The species identity was confirmed by DNA barcoding of all *B. terrestris* specimens, with sequences being identical to those of previously sequenced specimens of *B. terrestris* from this region (NCBI accession number: GU085204.1). Each of five males of *Bombus impatiens* was sourced from a distinct laboratory raised colony, which had been founded by field caught queens collected in Central Illinois, United States (GPS coordinates: latitude 40.657011; longitude -88.873755), in the spring of 2017. DNA was extracted from each bumblebee specimen using the Blood & Cell Culture DNA Mini Kit (Qiagen).

### Identification of TRs in the *B. terrestris* genome

Each of the 18 chromosome sequences of *B. terrestris* was uploaded to the Tandem Repeats Database (TRDB) (Gelfand et al. 2007). First, the sequence of each chromosome was analyzed using Tandem Repeats Finder (TRF) using default parameters: 2, 7, 7, 50 (match, mismatch, indels, minimal alignment score) (Benson 1999). As the bumblebee genome is AT-rich (~ 63%), poly A/T or AT/TA dinucleotides can occur by chance. Thus, to decrease the false positive rate of TR identification, TRs with repeat unit lengths of less than 2 or array lengths of less than 30 bp were discarded. Finally, redundant TRs reported for the same loci were excluded using the Redundancy Elimination tool at TRDB. For redundancy elimination, if TRs overlapped by more than 50% of their length, the repeat with the longer array was retained, or in the case of ties, the repeat with the shorter repeat unit length was retained. Manual correction was carried out when necessary.

### Characterizing the molecular features of TRs

The molecular features of TRs in *B. terrestris*, including repeat unit and repeat unit length distribution, TR array length distribution and genomic locations, were analyzed using the set of non-redundant TRs obtained from the above step by using a set of in-house Perl scripts, which are available at GitHub (https://github.com/suncheng781120/Tandem-repeat-analysis).

### Mining variable-length TRs between *B. terrestris* and *B. impatiens*

The sequence of each TR array, along with 100 bp of upstream and downstream flanking sequence, was extracted from the soft-masked *B. terrestris* genomic sequences (GCF_000214255.1). If there were continuous lower-case letters longer than 10 bp in either flanking sequence, indicating that the TR may reside in a repetitive region, the TR locus was excluded from further analysis. The sequences of the remaining TR loci, along with their 100 bp flanking regions, were used as queries in BLASTn searches against the genomic sequence of *B. impatiens*, with an e-value cutoff of 1e-10. For each query, we retained the best hit (based on e-value) that included both the TR array sequence and more than 95 bp of flanking sequences on both sides (because these hits likely represent the query’s orthologous locus in the *B. impatiens* genome). Finally, the pairwise alignments between the sequences of the TR arrays in *B. terrestris* and their best hits in *B. impatiens* were parsed to check if sequence length variation was observed within the TR array.

### Identification of TRs contributing to coding sequence variation

The coordinates of the identified variable-length TRs from the above step were used to search against the genome annotation of *B. terrestris* (downloaded from GenBank, see above) to identify those that resided in predicted coding DNA sequence (CDS). Then, whenever one variable-length TR was found in the coding sequence of one *B. terrestris* gene, the full-length protein sequence encoded by this *B. terrestris* gene was used as a query in a BLASTp search against the protein database of *B. impatiens* to find the best hit from *B. impatiens*. Finally, based on the pairwise alignments between the protein sequences of the query and its best hit, we checked for amino acid sequence variation caused by the variable-length TR (e.g., if one or more amino acid residues were added or deleted from one of the bumblebee species). If there was variation in the amino acid sequence, the variable-length TR was considered to contribute to bumblebee coding sequence variation.

### PCR amplification of identified variable TRs in coding sequences

The sequences of identified variable-length TRs residing in coding sequences, along with 200 bp of flanking sequences, were extracted from the genomic sequence of *B. terrestris*, and PCR primers were designed using Primer 3 (Untergasser et al. 2012). Then, with primers spanning the variable-length TRs, PCR was used to amplify genomic DNA samples extracted from *B. terrestris* and *B. impatiens* specimens (detailed PCR primer information is available in Supplementary file 1).

A 15 µL reaction mixture composed of 50 ng of template DNA, 0.3 µL of 10 mM each deoxynucleotide triphosphate (dNTP), 0.4 units of *Taq* DNA polymerase (Sangon Biotech, Shanghai, China), 1.5 µL of 10× PCR buffer with Mg^2+^, and 1.2 µL of 10 µmol/L forward and reverse PCR primers was prepared. Amplification was carried out using the following reaction conditions: initial denaturation at 94°C for 5 min, followed by 35 cycles of 30 s at 94°C, 30 s at 56°C, and 30 s at 72°C, with a final extension at 72°C for 10 min. 3 µL of PCR products were separated on 8% polyacrylamide denaturing gels, and the bands were revealed by silver-staining (Panaud et al. 1996).

### Functional classification of genes containing variable TRs

We used the predicted protein sequences of *B. terrestris* genes containing variable-length TRs as queries to do local BLASTp against the downloaded Swiss-Prot database (http://www.uniprot.org/uniprot/, last accessed on September 1, 2016), with an e-value cutoff of 1e-10. The UniProt accession of the best hit was used to represent this gene. The collected UniProt accessions were uploaded onto the PANTHER server (http://pantherdb.org/) and classified by the PANTHER system (Mi et al. 2013). If a TR-containing gene did not get a significant hit in the Swiss-Prot database or the obtained UniProt accession could not be mapped using PANTHER, we used the protein sequence encoded by the *B. terrestris* gene as query to search against the PANTHER library Version 12.0 (http://pantherdb.org/) with default settings to get a UniProt accession, which could be recognized by the PANTHER system, to represent the *B. terrestris* gene.

## Results

### The identification of TRs in bumblebee genome

We used the 18 chromosome sequences of *Bombus terrestris* (Sadd et al. 2015) as a reference in order to identify TRs in the bumblebee genome using the Tandem Repeats Finder algorithm (Benson 1999). After redundancy elimination (see Methods), a total of 26,595 TRs were identified. Our method focuses only on TRs that could be assembled and anchored on chromosomes (e.g., excluding those in highly-repetitive telomeric regions), which will underestimate the total number of TRs. Therefore, we do not intend to compare the absolute abundance of TRs in *B. terrestris* with that of related taxa.

### Molecular features of TRs in bumblebee

The distribution of repeat unit lengths of TRs in the bumblebee genome is summarized in Figure 1A. In general, the number of TR loci detected decreases with increasing repeat unit length. However, there are exceptions: two peaks occur at repeat unit lengths of 12 and 15 nt. The top ten most abundant repeat unit sequences, all either dinucleotide or trinucleotide, were quantified (Figure 1B), with the repeat unit “AG” as the most abundant in the bumblebee genome.

**Fig. 1.**
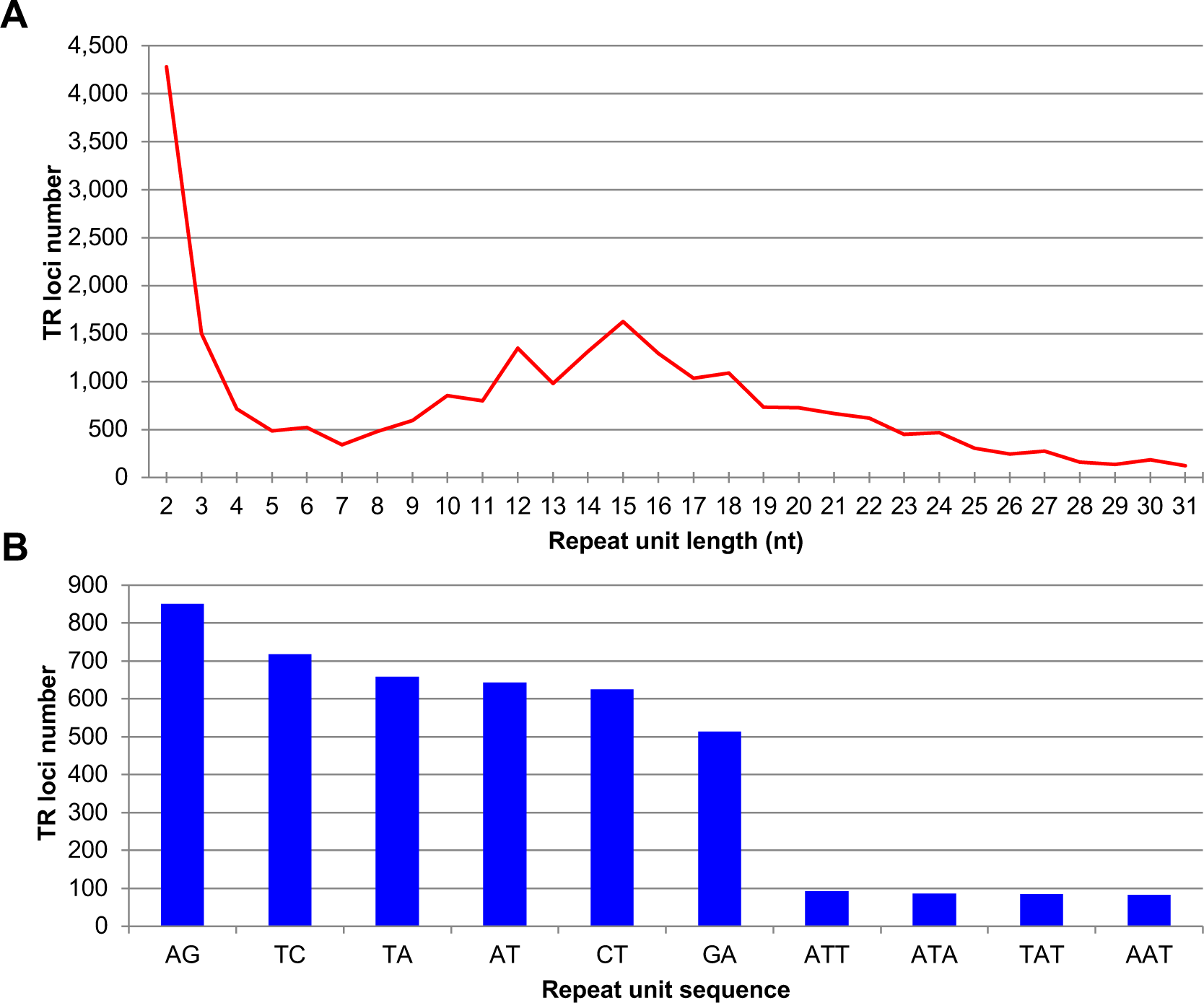
Repeat unit features for identified bumblebee TRs. (A) Repeat unit length distribution of TRs. Only repeat unit lengths, at which there are more than 100 TR loci, are shown. (B) The top 10 most abundant repeat unit sequences.

Most of the TR loci in the bumblebee genome identified by our methods are relatively short and 90% of TR loci have a length that is equal to or shorter than 111 base pairs (bps) (Figure 2A). To characterize the genome-wide distribution of TRs, the coordinates of TR loci were compared with the genome annotation for *B. terrestris*. Our results indicate that 66.7% (17,739 out of 26,595) of TRs in bumblebee genome were located within predicted genes (Figure 2B).

**Fig. 2.**
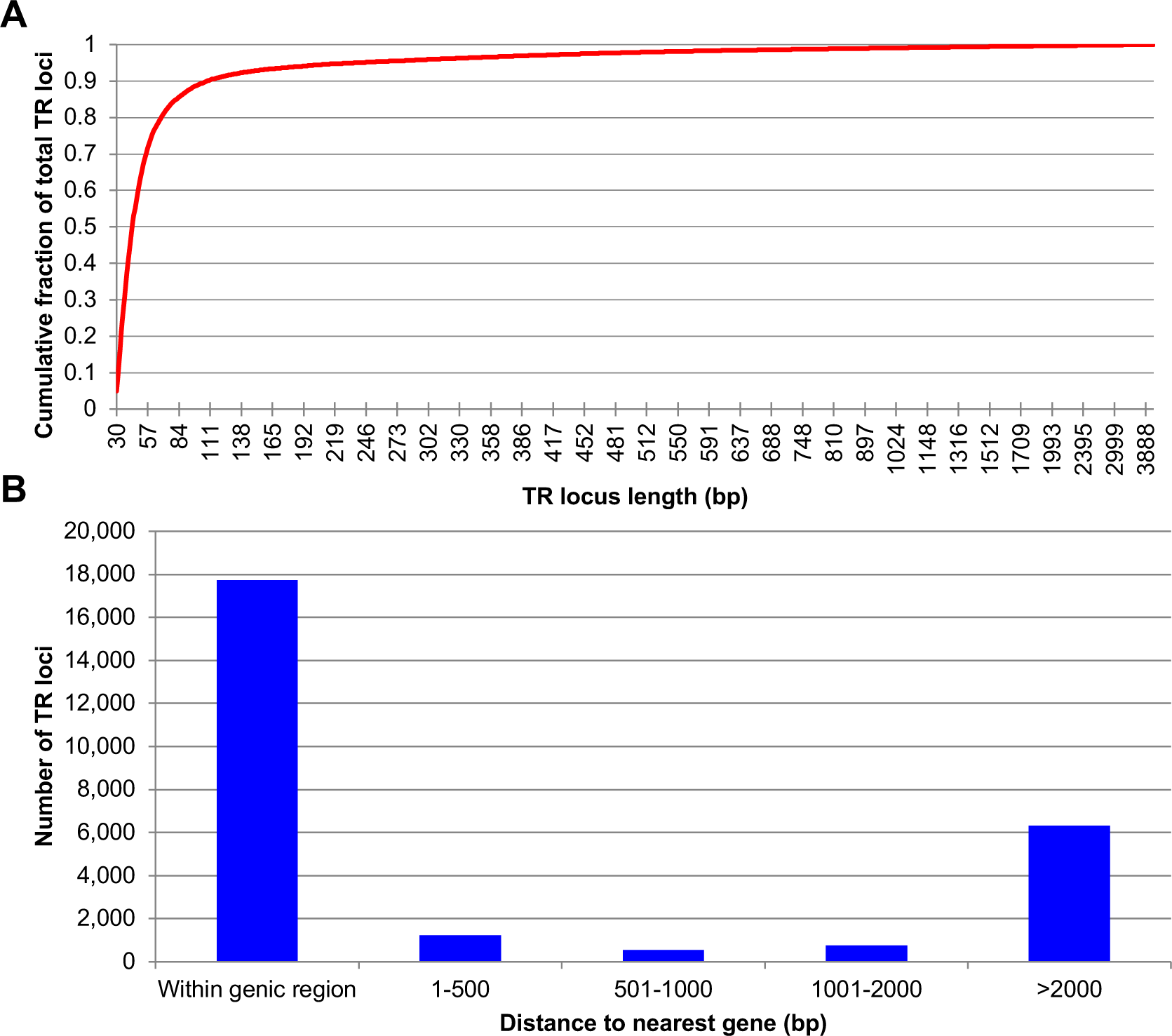
Distribution features for bumblebee TR loci. (A) TR locus length distribution. (B) The distance between TRs and predicted genes. As shown in the figure, a majority of TRs reside within genes.

### TRs contribute to genetic variation in bumblebees

To understand the contribution of TRs to genetic variation in bumblebees, TRs identified in the non-repetitive regions of the *B. terrestris* genome were used as queries to find their orthologous loci in another sequenced bumblebee genome, *B. impatiens*. Based on the pairwise alignments between the TR array sequences from the two bumblebee species, we identified variable TRs between them (see Methods). A total of 2,862 TRs were located within the non-repetitive regions of the *B. terrestris* genome, and, relative to *B. impatiens*, 1,137 of them are variable-length TRs (Supplementary file 2).

To understand if there are certain repeat unit lengths of TRs that are most likely to be array size variable between the two bumblebees, we calculated the ratio between the number of TRs showing variability in length between the two species and the number of TRs that do not exhibit variability in length for each repeat unit length, and plotted the ratio against the repeat unit length of TRs (Figure 3). Generally, TRs with repeat unit lengths ranging from 2 to 10 bp are more likely to be array size variable than longer TRs (Figure 3).

**Fig. 3.**
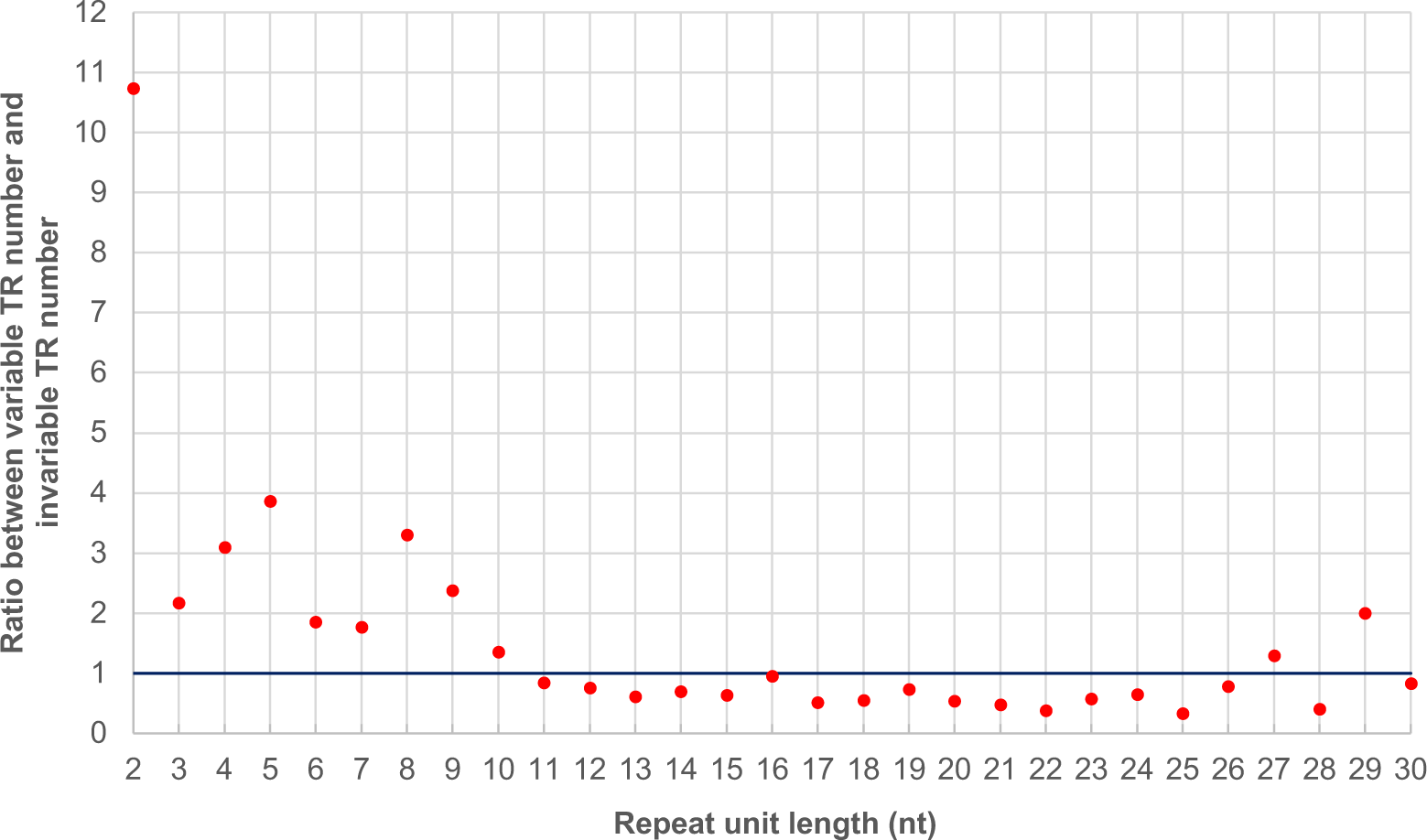
The relationship between repeat unit length and observed mutation probability for bumblebee TRs. The ratio between the number of TRs showing length variability and the number of TRs that do not exhibit variability in length was plotted against the repeat unit length of TRs.

### TRs contribute to coding sequence variation in bumblebees

To identify TRs generating coding sequence variation in bumblebees, we compared the genomic coordinates of the 1,137 variable TRs identified from the above step with those annotated as coding sequence (CDS) in the *B. terrestris* genome. We constructed pairwise alignments between protein sequences containing variable-length TRs to identify TRs generating protein sequence length variation between the two bumblebee species (see Methods). Based on this analysis, 101 of the 1,137 variable TRs were responsible for coding sequence variation (Supplementary file 3) and corresponded to protein sequence length variation (Supplementary file 4).

In Figure 4, we show one example of a TR generating coding sequence variation; the focal TR, which resides within a gene encoding a nuclear receptor corepressor, has a repeat unit of CAG (encodes glutamine), and there are five more repeat units in *B. terrestris* than in *B. impatiens* (Figure 4A). As a result, there are five more glutamine residues (represented by Q in the one-letter code) in the protein sequence encoded by the TR-containing gene in *B. terrestris* than in *B. impatiens* (Figure 4B).

**Fig. 4.**
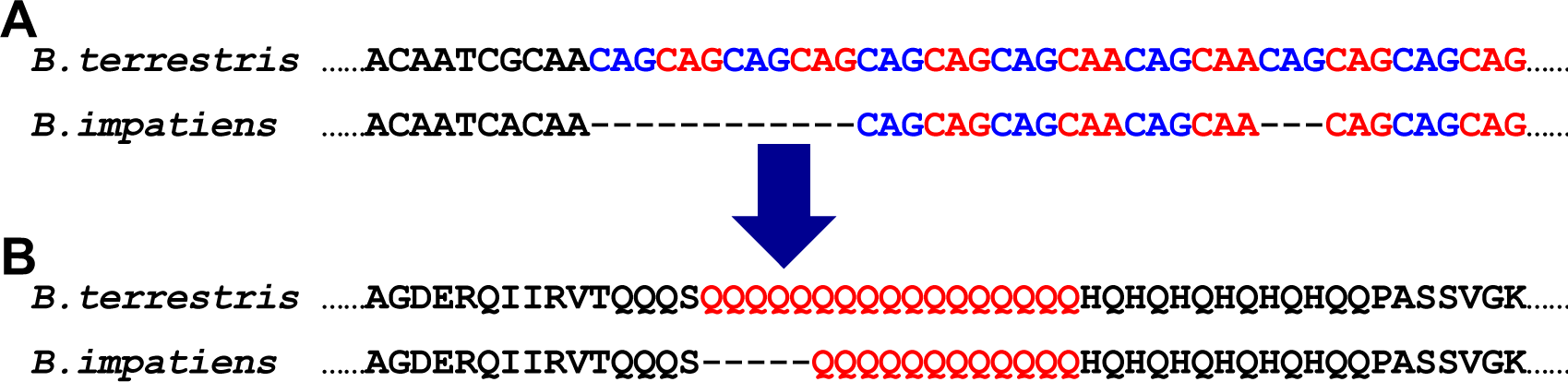
An example of TRs contributing to bumblebee coding sequence variation. (A) Pairwise alignments of TR arrays between *B. terrestris* and *B. impatiens*. Colored letters indicate TR array sequences, while black letters show their flanking sequences. The TR array has a repeat unit of CAG, and there are five more repeat units in *B. terrestris* than in *B. impatiens*. The coordinate for the variable TR is NC_015770.1:2190704-2190753 in *B. terrestris*. (B) Pairwise alignments of protein sequences encoded by genes containing the variable TR. Colored letters indicate TR array sequences, while black letters show their flanking sequences. There are five more glutamine residues (Q) in *B. terrestris* than in *B. impatiens*. Genes containing this variable TR encode nuclear receptor corepressor (protein IDs are XP_012166765.1 and XP_012249688.1 in *B. terrestris* and *B. impatiens*, respectively).

To further confirm that TRs contribute to coding sequence variation in bumblebees, we designed PCR primers that span the identified variable TRs in coding sequences and used them to amplify the genomic DNA extracted from 3 unrelated specimens of *B. terrestris* and 5 unrelated specimens of *B. impatiens* (Figure 5A). Our results (summarized in Table 1, with details available in Supplementary file 1) indicate that 19 of the 29 TR loci amplified exhibit interspecific length variation between *B. terrestris* and *B. impatiens*, with no length variation within species (denoted as Fixed variation). Eight of the 29 TR loci showed intraspecific length variation within at least one species, but the distributions of lengths in the two species do not overlap (denoted as Variation within species). Two of the 29 TR loci show trans-species variation, with overlapping distributions of length in the two species (denoted as Not fixed). Examples of the PCR amplification results revealing inter- and intraspecific variation of TRs in coding sequences can be seen in Figure 5 B and C, respectively. Altogether, our results suggest that TRs contribute to coding sequence variation in bumblebees.

**Fig. 5.**
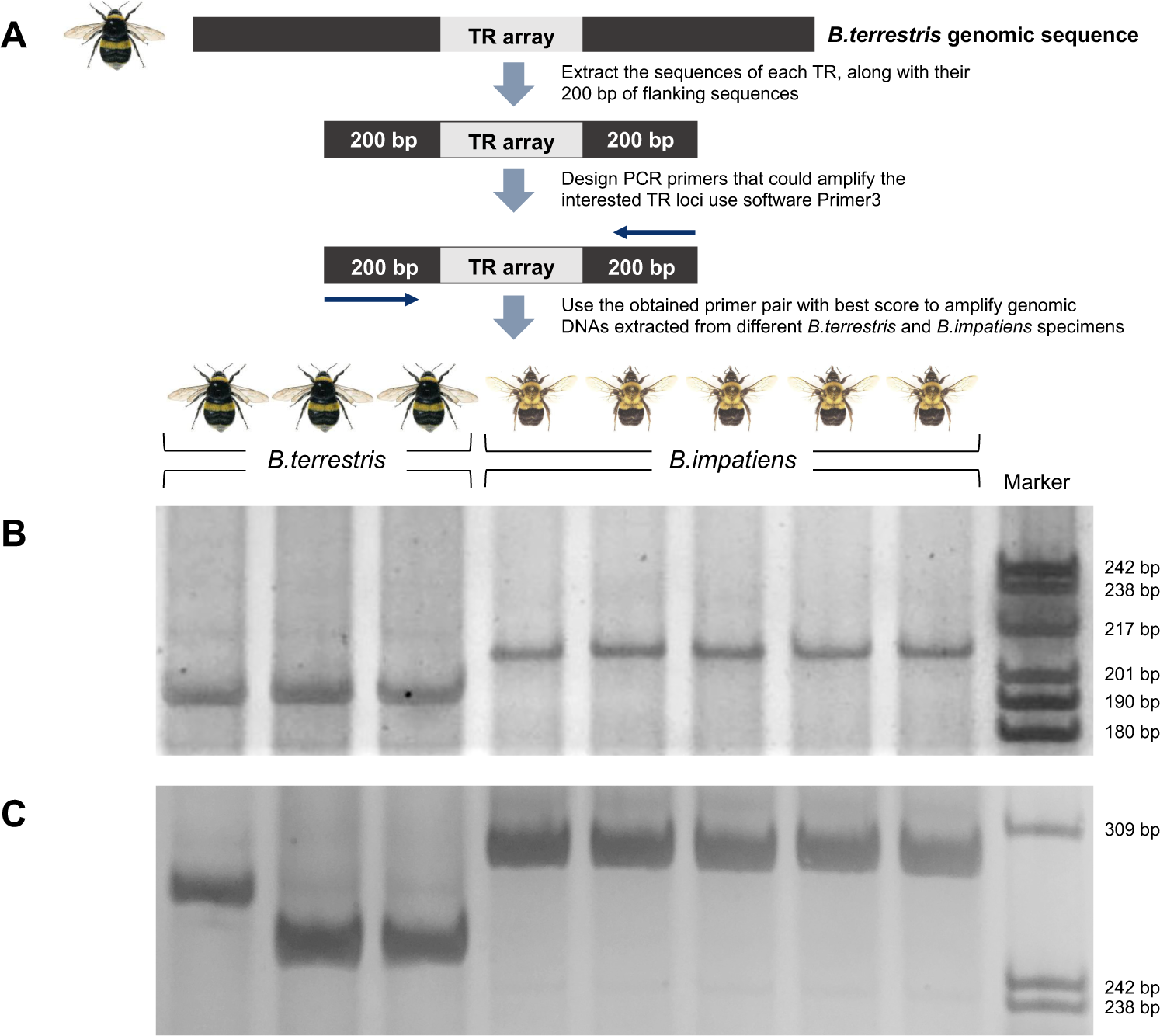
PCR amplification of variable-length TRs residing in coding sequences in specimens of *B. terrestris* and *B. impatiens*. (A) A schematic showing the principle of primer design and PCR amplification. (B) PCR amplification of the variable-length TRs residing in the gene that encodes FERM, RhoGEF and pleckstrin domain-containing protein (protein ID: XP_012169724.1 for *B. terrestris*). This figure indicates that, for the given TR locus, there is fixed length variation between the two species. (C) PCR amplification of the variable-length TRs residing in the gene that encodes hexamerin (protein ID: XP_012169664.1 for *B. terrestris*). This figure indicates that, for the given TR locus, there is length variation within species.

**Table 1.**
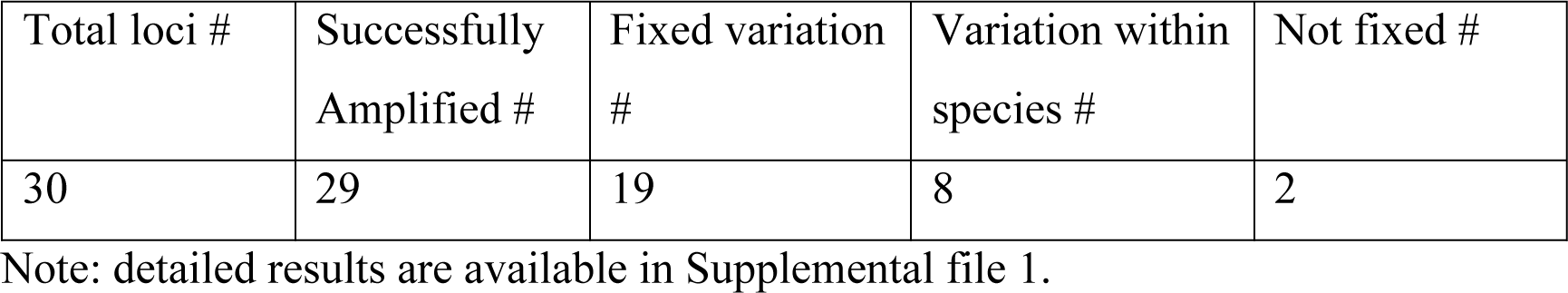
The summary for the PCR amplification of TR loci in coding sequences

We further examined the repeat unit length of the 101 variable-length TRs found in coding sequences. We observed 35 of them have a repeat unit length of 3, with all the other variable TRs having a repeat unit length of multiples of three (Supplementary file 5). This finding is consistent with previous research in other species, which indicates that selection should favor or tolerate mutations that avoid high impact frameshift mutations (Legendre et al. 2007; Mularoni et al. 2010; Richard and Dujon 2006; Young et al. 2000).

### Protein-coding gene sequence variation driven by TRs in bumblebees

The identified 101 variable TRs that contribute to coding sequence variation in the sequenced bumblebees are found in 85 protein-coding genes. We performed a functional classification using PANTHER, from which 74 of them could be functionally classified. Over half of the classified genes (26 out of the 48 genes that could be assigned a molecular function) are involved in binding, which is defined as the selective, non-covalent, often stoichiometric, interaction of a molecule with one or more specific sites on another molecule (Figure 6A). The second most frequent molecular function is catalytic activity, with 15 genes falling in this category. Other molecular functions of classified genes include structural molecular activity, receptor activity, and transporter activity (Figure 6A).

**Fig. 6.**
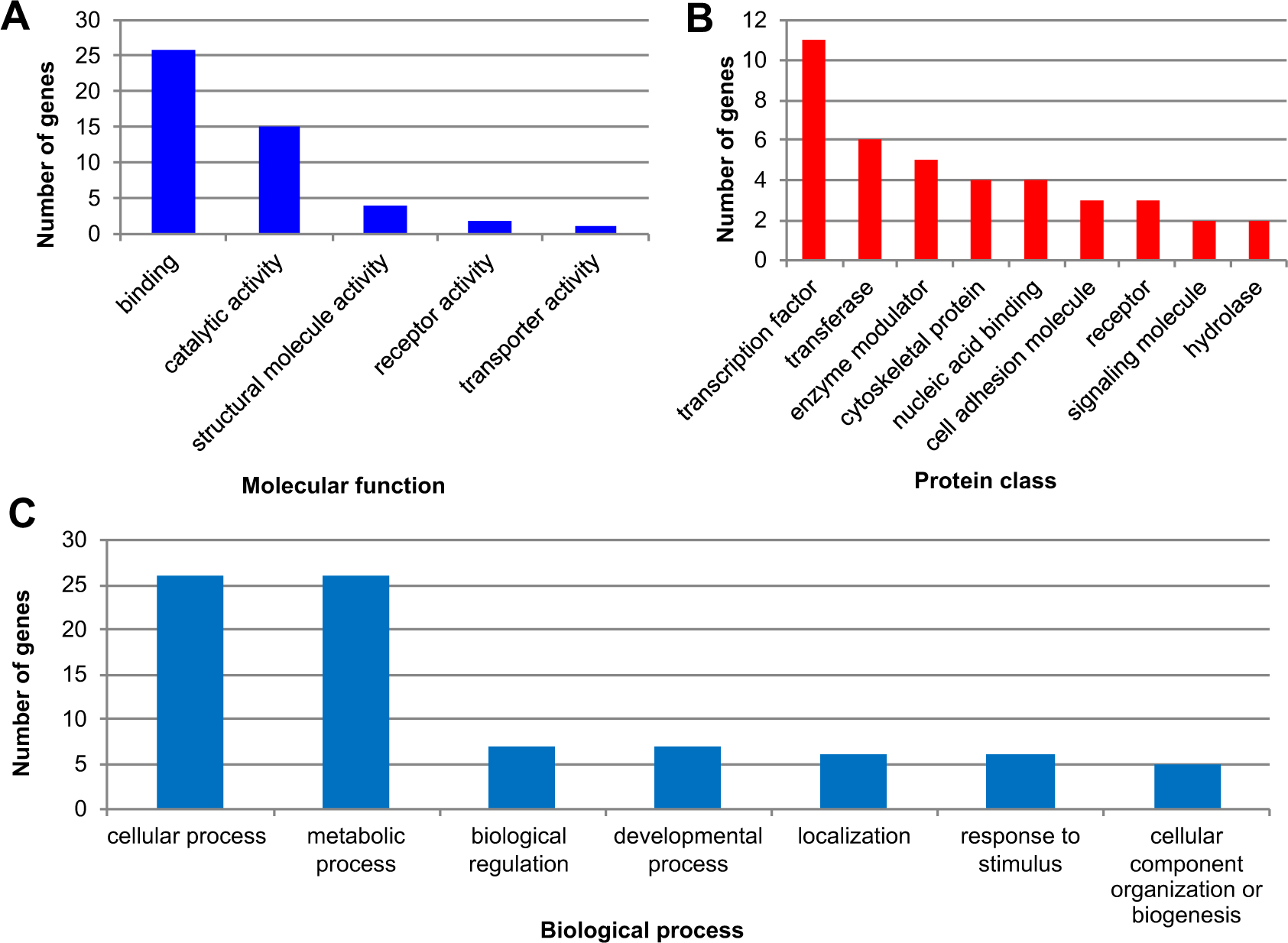
Functional classification of genes that include variable-length TRs. (A) The number of genes classified in each molecular function category. (B) The number of genes classified in each protein class. The gene number shown in the nucleic acid binding category excludes transcription factors. (C) Biological processes that genes including variable-length TRs are involved in.

Proteins encoded by those genes containing variable-length TRs were assigned to 18 protein class categories, and the top 9 categories (categories having two or more genes) are shown in Figure 6B. The most frequent protein class category represented is transcription factors, and a total of 11 genes were found to encode them (Figure 6B). Based on the available databases that curate transcription factors in the genome of *B. terrestris*, the KEGG BRITE database (http://www.genome.jp/kegg/brite.html) identifies 271 genes as eukaryotic transcription factors (last updated on June 8, 2018), and the Regulator database (http://www.bioinformatics.org/regulator) returns 529 metazoan transcription factors (including 76 potential transcription factors, last accessed on July 5, 2018,). Thus, based on the current assignment of the Regulator database, there are 529 transcription factors in *B. terrestris* among the ~13,000 protein-coding genes in the genome (Kapheim et al. 2015). While 4.06% (529 out of 13,000) of bumblebee genes encode transcription factors, 12.94% (11 out of 85) of the classified genes containing variable-length TRs are transcription factors, which represents a >3-fold overrepresentation in this category. Other identified protein class categories include transferase and enzyme modulators (Figure 6B).

Bumblebee genes where coding sequences contained variable-length TRs are involved in a variety of biological processes (Figure 6C). The most frequent biological process categories are cellular and metabolic processes, each with 26 classified gene. Other biological processes represented include biological regulation, developmental processes, and response to stimulus. Genes containing variable-length TRs were involved in eight known pathways, namely, Wnt signaling, Nicotinic acetylcholine receptor signaling pathway, Apoptosis signaling pathway, Alzheimer disease-presenilin pathway, 5HT2 type receptor mediated signaling, p38 MAPK pathway, Heterotrimeric G-protein signaling pathway, and Huntington’s disease pathway. Interestingly, one bumblebee gene where the coding sequence contains variable-length TRs has the same tri-nucleotide repeat expansion(CAG) as that which causes Huntington’s disease in humans (Figure 4), and was determined to be associated with the Huntington’s disease pathway by PANTHER.

## Discussion

Genomic tandem repeats (TRs) are widespread in diverse species, where they represent highly dynamic regions of mutation, which can facilitate the rapid evolution of coding and regulatory sequences (Gemayel et al. 2010). However, to date, little is known about TRs in bumblebees, despite their importance as pollinator species and their wide range of habitats (Sakagami 1976; Williams 1985). The present study represents the first systematic analysis of TRs in bumblebees. Based on our search criteria, there are over twenty-six thousand TRs in *B. terrestris* genome, and 1,137 of which are polymorphic when compared to a closely related species, *B. impatiens*. Our TR identification method may underestimate the true number of TRs in bumblebees. Although it will find micro- and minisatellite sequences, it is too stringent to identify larger satellite sequences. Also, our method likely underestimates the true number of variable-length TRs among species of bumblebee because we only included TRs in non-repetitive regions (2,862) for subsequent analysis (see Methods). Furthermore, variable-length TRs were identified based on a comparison of only two bumblebee species. There are 38 subgenera of bumblebees, and *B. terrestris* and *B. impatiens* are only representative of two (Cameron et al. 2010; Hines 2008). Therefore, our results represent a conservative estimate of the effect of TRs on bumblebee genetic variation, and the true amount of sequence variation contributed by TRs is likely much greater.

Because genetic variation is an essential starting point for adaptation to new environments (Barrett and Schluter 2008; Lande and Shannon 1996; Paaby and Rockman 2014), we postulate TRs may contribute to adaptation of bumblebees across the many niches in which they are found. With threats to bumblebees of upmost concern given recent population declines (Cameron et al. 2011), TRs may also determine susceptibility and evolutionary responses to proposed environmental stresses (Goulson et al. 2015).

Interestingly, in this study, we find evidence for changes in protein-coding sequence due to variation in TRs, and the frequency of such changes are most frequently observed in proteins known to influence gene expression. Both changes in protein sequences and changes in gene expression could drive adaptation, although the relative importance of these two molecular mechanisms has long been controversial (Fondon and Garner 2004; Fraser 2013; Hancock et al. 2011; King and Wilson 1975; Wray 2007). To understand the possible molecular mechanisms facilitating adaptation in bumblebees through TRs, we focus on changes in protein sequences rather than changes in gene expression, because even *cis*-regulatory sequences, which are directly related to changes in gene expression (Wray 2007), have not been extensively annotated yet in the bumblebee genomes. In this study, we searched for TRs that generate coding sequence variation, which in turn produce proteins of varying lengths (Supplementary file 4). In terms of the protein-coding changes we observed, for the 101 variable-length TRs identified, all the repeat units have a length of multiples of three (Supplementary file 5), which is consistent with findings in other species suggesting that natural selection may favor mutations that avoid frame-shifts (Legendre et al. 2007; Mularoni et al. 2010; Richard and Dujon 2006; Young et al. 2000). Mutations in TRs altering the length of protein sequences without introducing frame-shifts have the potential to majorly increase the functional diversity of host genes (Caburet et al. 2005; Fondon and Garner 2004; Gemayel et al. 2010; Radó-Trilla et al. 2015; Verstrepen et al. 2005). Our functional classification, however, further revealed that the most frequent protein class category exhibiting variable-length TRs is transcription factors, with a total of 11 genes (Figure 6B, Supplementary file 5), which is a >3-fold overrepresentation relative to the expectation (see Results). Changes to the coding sequence of transcription factors could change their three-dimensional structure, target binding site, specificity, and their ability to recruit other transcription factors. Most importantly, changes in transcription factors could lead to modified transcription levels of genes at many other loci in the genome, in contrast to protein-coding changes in structural or signaling proteins which only affect the protein in which they occur.

Organisms can adapt to new environments by regulating gene expression at multiple stages of mRNA biogenesis, a process governed by many different proteins, such as transcription factors, chromatin-remodeling factors, signaling molecules, and receptors (De Nadal et al. 2011; Kadonaga 2004). The second and the third most frequent protein class categories, transferases and enzyme modulators, respectively, are also involved in gene expression regulation (Figure 6B). We checked all these protein class categories manually, and identified a total of 34 genes (out of the 39 genes that could be assigned to a protein class by PANTHER) that are involved in regulating gene expression (Supplementary file 5). Altogether, our results indicate that TRs in bumblebee drive potentially functional variability at loci involved in gene expression regulation and other biological functions. As a result, length variation of TRs may facilitate the adaptation of bumblebees through diversifying bumblebee proteins, particularly those which regulate gene expression, as has been previously hypothesized (Fraser 2013; King and Wilson 1975; Wray 2007).

## Conclusions

In the present study, we performed a comprehensive investigation of TRs in bumblebees. Our results indicate that TRs are prevalent in sequenced bumblebee genomes, and a majority of those identified reside within genic regions. We found out that TRs represent a significant source of genetic variation in bumblebees. They promote coding sequence variation and influence the functional diversity of bumblebee genes. The functional roles of genes whose coding sequences contain variable-length TRs were analyzed, and our results indicate that a majority of those genes are related to transcriptional regulation. Given the importance of gene expression changes for adaptation, our observation that loci encoding transcription factors are enriched for variable-length TRs may suggest an important role for expanded repeats in the evolution of bumblebees.

### Data deposition

All data generated or analyzed during this study are included in this published article (and its Supplementary files).

## Acknowledgements

This work was supported by the Science and Technology Innovation Project of Chinese Academy of Agricultural Sciences [CAAS-ASTIP-2017-IAR] and the Elite Youth Program of Chinese Academy of Agricultural Sciences [to CS]; National Science Foundation [MCB-1150213] funding [to SS]; and a National Science Foundation [IOS 16-54028] and an Illinois State University Pre-tenure Faculty Research Initiative Grant [to BS]. We thank Dr. Jiaxing Huang and Jiandong An (Institute of Apicultural Research, Chinese Academy of Agricultural Sciences) for providing excellent assistance with bumblebee collection in China.

## Supplemental material Legends

**Supplementary file 1.** This file contains detailed information for the PCR amplification of TR loci in coding sequences, which includes TR loci coordinates, primer sequences, PCR reaction conditions and amplification results etc.

**Supplementary file 2.** This file shows the pairwise alignments of the identified 1,137 variable-length TRs between *B. terrestris* and *B. impatiens*.

**Supplementary file 3.** This file shows the pairwise alignments of the identified 101 variable-length TRs in coding sequences between *B. terrestris* and *B. impatiens*.

**Supplementary file 4.** This file shows the pairwise alignments of proteins sequences encoded by genes containing variable-length TRs between *B. terrestris* and *B. impatiens*.

**Supplementary file 5.** This file shows the coordinates and repeat unit length for the 101 variable-length TRs in coding sequences. Proteins encoded by genes harboring those variable-length TRs and their corresponding protein class categories, if related to transcription regulation, are also shown.

